# Introduction of cytosine-5 DNA methylation sensitizes cells to oxidative damage

**DOI:** 10.1101/2024.09.13.612259

**Authors:** J. Krwawicz, C.J. Sheeba, K. Hains, T. McMahon, Y. Zhang, S. Kriaucionis, P. Sarkies

**Affiliations:** Department of Biochemistry, University of Oxford, Oxford, UK; London Institute of Medical Sciences, Imperial College London, Du Cane Road, London, UK; Ludwig Institute for Cancer Research, University of Oxford, Oxford, UK

## Abstract

DNA methylation at the 5 position of cytosine (5mC) is an ancient epigenetic mark in eukaryotes. The levels of total 5mC vary enormously between different species, and the DNA methyltransferases that introduce 5mC have been repeatedly lost in several independent lineages. DNA methyltransferases are a threat to genomic stability due to the increased mutagenicity of 5mC bases and the propensity of DNA methyltransferases themselves to introduce DNA alkylation damage as an off-target effect. However, whether alkylation damage explains why 5mC is frequently lost in evolution is unclear. Here we tested the fitness consequences of DNA methyltransferase-induced alkylation damage by introducing a eukaryotic-like 5mC system into *E. coli*. We showed that introducing 5mC genome-wide leads to increased sensitivity to alkylating agents, which is strongly enhanced by removal of the 3mC repair enzyme AlkB. Unexpectedly, we discovered that 5mC introduction led to increased sensitivity to oxidative stress. We showed that this is due to increased formation of reactive oxygen in the presence of 5mC. We determined that reactive oxygen species led to non-enzymatic oxidation of 5mC, producing modified cytosines such as 5fC that are recognised as DNA base damage in *E. coli*. Overall, our work identifies increased sensitivity to oxidative stress, as well as alkylating agents, as a negative consequence of genome-wide 5mC. Oxidative stress is frequently encountered by organisms in their environment, thus offering a plausible reason for total loss of 5mC in some species.

## Introduction

Methylation of the 5^th^ position of cytosine within DNA (5mC) is found in many species drawn from all domains of life(Jurkowski & Jeltsch, 2011). In prokaryotes (bacteria and archaea) methylation is carried out by DNA methyltransferases (DNMTs) that typically have sequence specificity targeting them to 4-8bp sequence motifs. 5mC introduced in this manner contributes to restriction modification systems, which protect genomes against invading viruses, plasmids or other mobile genetic elements. In contrast, in eukaryotes, DNMTs that introduce 5mC have more limited sequence specificity, and are capable of acting genome-wide. Across eukaryotes, 5mC is commonly introduced at CG dinucleotides(de Mendoza et al., 2020). The symmetrical nature of this sequence context enables 5mC in the CG context to act epigenetically, because maintenance methyltransferases (such as DNMT1 in mammals and Met1 in plants) have evolved to recognise hemi-methylated DNA after DNA replication and introduce methylation onto the unmethylated strand(Law & Jacobsen, 2010). This means that 5mC in CG contexts can persist through cell division independent of the activity of the de novo methyltransferases (such as DNMT3A and B in mammals) that introduced them initially. DNA methylation in other sequence contexts is widespread in plants and is also found in some vertebrate genomes, particularly in non-dividing cells such as neurons(de Mendoza et al., 2020, 2021).

In eukaryotes, the functions of DNA methylation has not so far been linked to restriction modification systems analogous to bacteria. Instead, 5mC has roles in gene expression regulation(Sarkies, 2022b). These roles are complex, overlapping and not fully characterised, especially because they depend on genomic context. For example, in mammals DNA methylation at promoter regions is associated with transcriptional silencing during development(Weber et al., 2007), whereas methylation in gene bodies is linked to the presence of histone modifications associated with transcription, such as H3K36me3 and may contribute to noise suppression for example through suppression of spurious transcription (Baubec et al., 2015). In arthropods, 5mC is much less abundant genome-wide and is often confined to a small number of transcriptionally active housekeeping genes(Bewick et al., 2017; Lewis et al., 2020).

Eukaryotic 5mC methylation, and, in particular, epigenetic methylation in the CG context has an ancient origin. Nevertheless, DNA methylation is far from universal. Many species have lost DNA methylation altogether, including many commonly used as model organisms such as *Drosophila melanogaster*, *Caenorhabditis elegans*, *Saccharomyces cerevisiae and Schizosaccharomyces pombe*(Bewick et al., 2019; Feng et al., 2010; Keller et al., 2016; Zemach et al., 2010). As indicated above, the levels of 5mC vary enormously across different species, even if they possess methylation machinery(Bewick et al., 2017; Lewis et al., 2020). Often evolutionary transitions associated with either widespread increases in DNA methylation levels or complete loss have occurred very recently, suggesting ongoing processes selecting for or against DNA methylation in different species(de Mendoza, Hatleberg, et al., 2019; de Mendoza, Pflueger, et al., 2019). The reasons for such rapid diversification in methylation systems are still largely unknown.

One important possible explanation that has emerged is the propensity of 5mC to damage the genome. This is observed as an increased rate of mutations at methylated CpGs, when compared to unmodified contexts(Tomkova & Schuster-Böckler, 2018). In part this is due to the fact that cytosine deaminates to uracil whilst 5mC deaminates to thymine(Lindahl, 1996).Recognition of uracil as a foreign DNA base might be easier in cells than detecting a T-G mismatch, leading to an elevated rate of C to T mutations at 5mC(Schmutte et al., 1995; Tarantino et al., 2018). Potential evidence for this was first provided by the observation of depletion of CG dinucleotides in mammalian genomes relative to the expected proportion based on the total number of C and G nucleotides(Bird, 1980; Gardiner-Garden & Frommer, 1987). Subsequently, analysis of mutation rates in cancer quantified the excess C-T mutation rate(Alexandrov et al., 2013), leading to a more sophisticated understanding of its origin including the discovery that 5mC increases the rate of base pair misincorporation by replicative polymerases, particularly the leading strand DNA polymerase Polε (Tomkova et al., 2018, 2024).

In addition to increased mutation of 5mC nucleotides, we recently discovered an additional source of genome instability due to DNMT activity. We discovered that across eukaryotes DNMTs co-evolve with alkylation repair, in particular the enzyme ALKBH2, which tends to be present whenever DNMTs are present and absent when DNMTs have been lost(Rošić et al., 2018). In arthropods, the presence of ALKBH2 correlates with higher levels of 5mC genome wide(Lewis et al., 2020). ALKBH2 repairs the damaged base 3mC, and indeed we showed that DNMTs can introduce 3mC *in vitro* and *in vivo*(Dukatz et al., 2019; Rošić et al., 2018).

The mutagenic and genotoxic consequences of DNMT expression might be one reason why DNMTs, and 5mC could be selectively disadvantageous. This cost might be more important if the species is exposed to high levels of DNA damage in its environment, thus putting more strain on DNA repair mechanisms(Sarkies, 2022b). Here, we set out to explore this hypothesis to attempt to test directly the consequences of DNMT activity for sensitivity to genotoxic stress. In order to test this independent of functions of DNMT in gene regulation, we developed a system to introduce high levels of CG methylation into the model bacteria *E. coli*. We demonstrated that *E. coli* expressing DNMTs was hypersensitive to DNA alkylating agents that introduce 3mC into the DNA, and that this was exacerbated by loss of AlkB, the *E. coli* homologue of ALKBH2(Fedeles et al., 2015). Unexpectedly, we also discovered that DNMTs led to hypersensitivity to oxidative stress induction. We showed that DNMT expression leads to increased accumulation of reactive oxygen species, which is exacerbated by exogenous oxidising agents. We present biochemical and genetic evidence that the consequence of oxidative damage in genomes with 5mC is the emergence of 5hmC and 5fC - further oxidation products of 5mC, which results in toxicity. Taken together, our data points to an additional source of genotoxic stress beyond alkylation damage associated with DNMT activity, which could have particular relevance for explaining why some species have lost DNMTs and 5mC completely, and may also be important in the cellular consequences of DNMT activity in mammals.

## Results

### A system to introduce genome-wide CG methylation into E. coli

Studying the effect of DNMT activity on genotoxic stress sensitivity is complicated in eukaryotic cells because CG methylation is associated with gene expression so secondary effects on genotoxic stress due to expression changes are likely to be pervasive. We therefore sought to establish a system where we could introduce DNMT activity into cells and study its effects on genotoxic stress more straightforwardly. We took advantage of the highly active bacterial CG methyltransferase M.SssI(Darii et al., 2009). The catalytic residues of M.SssI are highly conserved with eukaryotic DNMTs (Figure 1A). We expressed M.SssI on a low copy number plasmid in a strain of *E. coli* lacking the McrABC restriction modification system (C2523). This resulted in robust induction of genome-wide 5mC in the CG content, as demonstrated by methylation sensitive restriction digest and mass spectrometry (Figure 1B,C). We also expressed the CG methyltransferase M.MpeI as an alternative model to test our conclusions(Wojciechowski et al., 2013). M.MpeI induced lower levels of 5mC genome-wide than M.SssI (Figure 1C). Together these two enzymes provided an opportunity to study the effects of genome-wide 5mC in the CG context on genotoxic stress.

**Figure 1.**
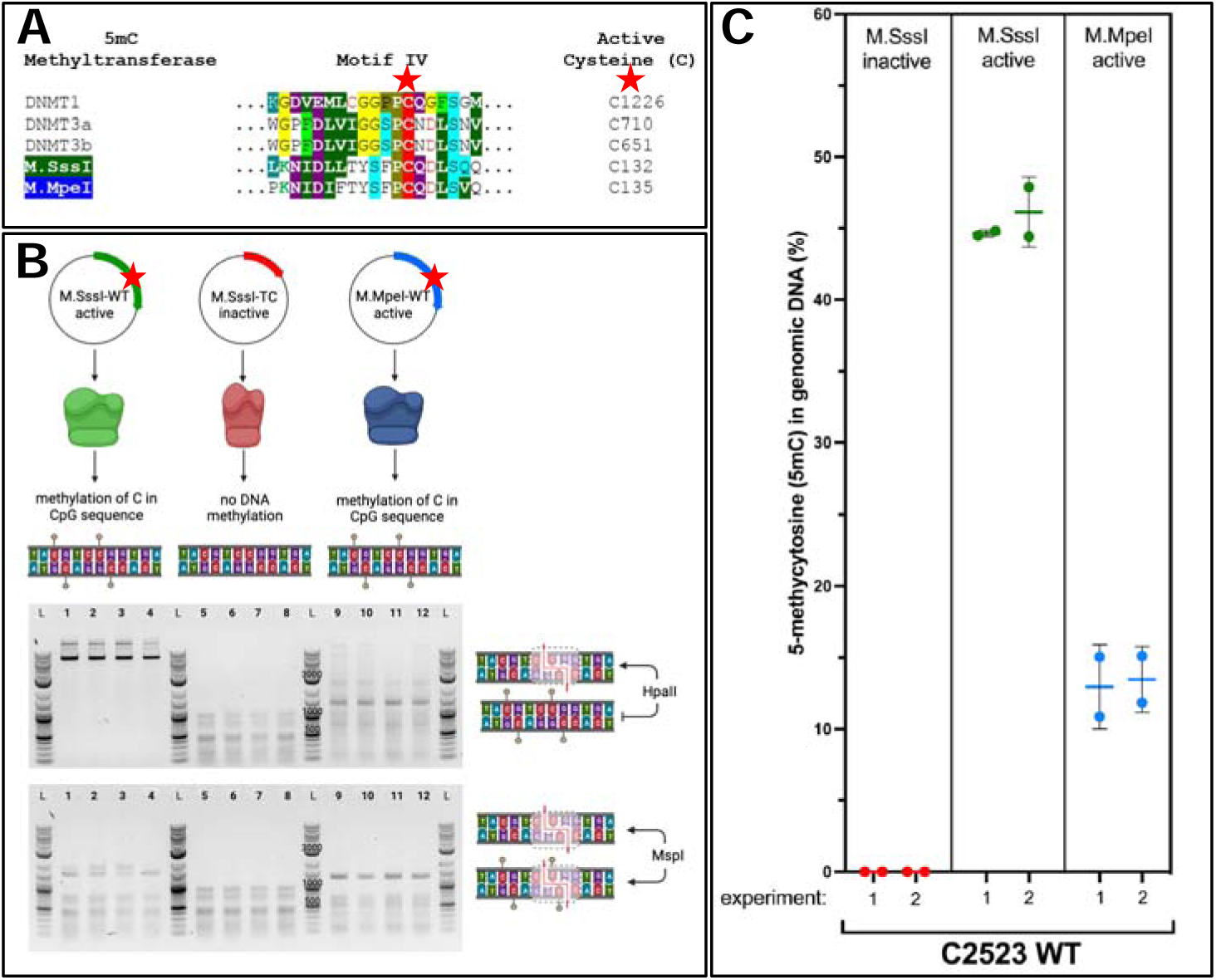
- Introduction of genome-wide CG methylation into *E. coli*. A – Plasmids encoding active 5mC DNA methyltransferases (M.SssI and M.MpeI) and inactive M.SssI were introduced into *Esherichia coli* strain B C2523 without active 5mC nuclease McrA to protect methylated DNA before degradation. B Methyl-sensitive restriction enzyme digest. HpaII is blocked by 5mC in the recognized C5mCGG palindrome and MspI can digest methylated C5mCGG and unmethylated CCGG sequences. C Genomic DNA was isolated from strains expressing active methyltransferases (M.SssI and M.MpeI) and inactive M.SssI, and global 5mC level was calculated as a percentage of total cytosine.

### DNMTs sensitize cells to alkylation damage

Our previous work demonstrated that DNMTs induce alkylation damage in the form of 3mC into genomic DNA(Rošić et al., 2018). To investigate the effect of this on genotoxic stress resistance we exposed cells expressing DNMT to the DNA alkylating agent Methyl-methan-sulfonate (MMS). MMS introduces a number of alkylation lesions into DNA including 3mC(Sedgwick, 2004). Cells expressing either M.SssI or M.MpeI showed increased sensitivity to MMS treatment compared to cells expressing inactive M.SssI (Figure 2), supporting the conclusion that the expression of DNMTs increased the levels of alkylation damage. To test this conclusion further we inactivated AlkB, the *E. coli* homologue of the eukaryotic ALKB family(Ougland et al., 2015). *E. coli* AlkB, similarly to ALKBH2, is able to repair 3mC damage in DNA(Trewick et al., 2002), although it should be noted that its substrate specificity is much wider, with the ability to act on RNA and protein as well as DNA(Fedeles et al., 2015). *alkB* mutants were hypersensitive to MMS treatment compared to WT C2523, consistent with previous studies of the *E. coli* AlkB enzyme(Dinglay et al., 2000; Sikora et al., 2010; Trewick et al., 2002). Expression of DNMTs strongly exacerbated this effect (DNMT p=6.4e-14 two-way Anova, Figure 2). Importantly, DNMT expression combined with deletion of AlkB had a greater effect on MMS sensitivity than would be expected from the effect of each individually (DNMT\**alkB* interaction p=0.0044 three-way Anova). In order to investigate the non-additivity between DNMT expression and *alkB* mutation further, we investigated the effect of MMS over a range of concentrations for the different strains (Supplemental Figure 1A). We quantified the non-additivity by comparing between the survival of alkB expressing DNMT to the predicted combined effect of either *alkB* mutation alone or DNMT expression alone (Supplemental Figure 1B). Significantly reduced survival than expected was observed, most notably at low concentrations of MMS, which could be due to the saturation of the effect at high concentrations of MMS for *alkB* mutants expressing DNMT, where extremely high levels of sensitivity were observed. The non-linear shape of the graph observed for WT cells expressing DNMTs further suggests that the ability of AlkB to repair the DNA is overwhelmed at high MMS concentrations even in the WT background. These results are consistent with the idea that AlkB repairs a form of DNA damage from MMS that is more prevalent when DNMT is expressed. This could be because DNMT induces 3mC, repaired by AlkB, and further 3mC is induced by MMS leading to much higher 3mC levels in the absence of AlkB activity. Alternatively, 3mC induction by DNMT may lead to increased levels of ssDNA, particularly in *alkB* mutants, which could increase the risk of further DNA damage by MMS exposure and heighten sensitivity. Either of these mechanisms are consistent with induction of 3mC by DNMT.

**Figure 2.**
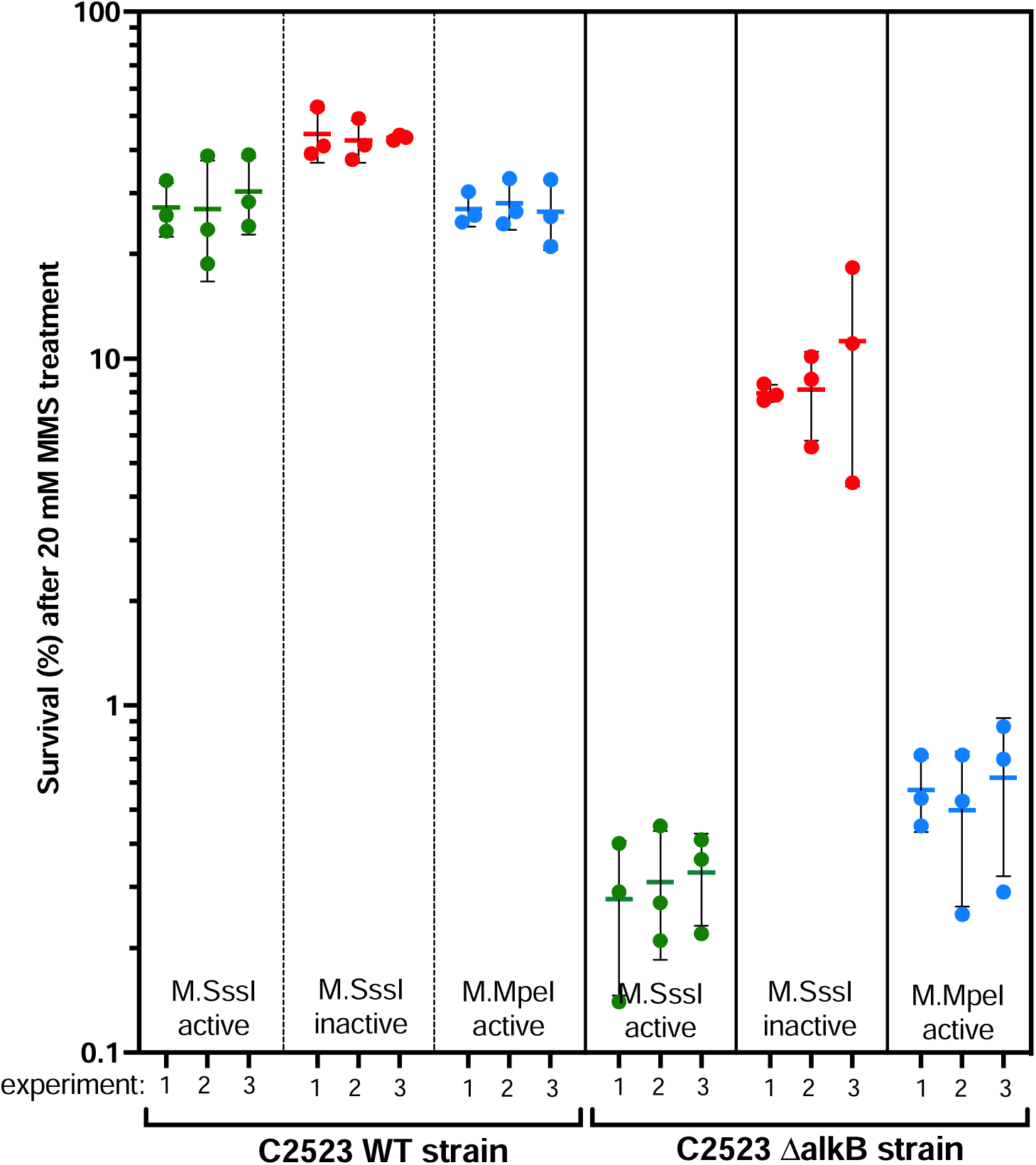
MMS-induced sensitivity in strains carrying active 5mC methyltransferases. WT strains and *alkB* mutants with M.SssI and M.MpeI methyltransferases were treated with 20 mM MMS. The y-axis shows the percentage survival relative to the survival under the same conditions without MMS. Each set of 3 points shows three separate clones derived from a single transformation with the indicated plasmid. Three separate transformations were performed for each plasmid.

One potential consequence of DNMT activity in inducing DNA damage might be increased mutagenesis. To test this we performed a rifampicin resistance mutagenesis assay, in the absence of MMS, to test whether DNMT induced damage was sufficient to lead to mutation rate increase. Mutation rate was increased by DNMT expression (p=1.6e-12; two way anova; Supplemental Figure 2) and *alkB* mutation (two way anova) separately (p<1e-16). Moreover, there was a significant interaction such that combined *alkB* mutation and DNMT expression led to a further increased mutation rate compared to the expectation from *alkB* mutation and DNMT expression separately (p = 7.9e-10; Supplemental Figure 2). Importantly, DNMT induction alone would be expected to lead to increased mutations due to cytosine deamination(Sarkies, 2022a); however, there is a synergistic effect on mutations when this is combined with loss of AlkB function in *alkB* mutants. This is consistent with 3mC induction by DNMTs which is repaired by AlkB in WT cells but leads to mutations in *alkB* mutant cells.

Overall these results indicate that the induction of DNA damage by DNMT expression has increases sensitivity to genotoxic stress in their environment and results in increased mutagenesis even in the absence of genotoxic stress. These effects indicate a substantial fitness cost for DNMT expression in *E. coli*.

### DNMT expression sensitizes cells to oxidative stress

We wondered whether the effect of DNMT expression was specific to DNA alkylation damage sensitivity. We therefore subjected DNMT expressing cells to other genotoxic stresses and tested their sensitivity. WT C2523 cells and C2523 expressing DNMT were equally sensitive to cisplatin (Supplemental Figure 3), which introduces intra and interstrand cross-links into the DNA(Dasari & Bernard Tchounwou, 2014). However, DNMTs demonstrated a marked sensitivity to induction of oxidative stress by exposure to H_2_O_2_ (Supplemental Figure 3; Figure 3). Genotoxic stress associated with H_2_O_2_ treatment is due to production of reactive oxygen species which generate a number of lesions in DNA(Svilar et al., 2011). Oxidised purines in particular are excised by the base excision repair glycosylase Fpg, also known as MutM, homologous to the mammalian NEIL family(Hwang et al., 2025). We deleted Fpg in *E. coli* and investigated sensitivity to oxidative stress. Fpg deletion alone had little effect, presumably due to redundancy across DNA repair pathways (Figure 3). However, induction of active DNMT expression through M.SssI led to a strongly exacerbated sensitivity to H_2_O_2_ treatment (p<1e-16 Two way anova; Figure 3). The combined effect of expression of DNMT and mutation of Fpg was greater than the sum of the individual effects (Fpg*DNMT interaction p=0.032; Figure 3). Together this suggested that DNMT expression increased sensitivity to oxidative stress-induced DNA damage.

**Figure 3.**
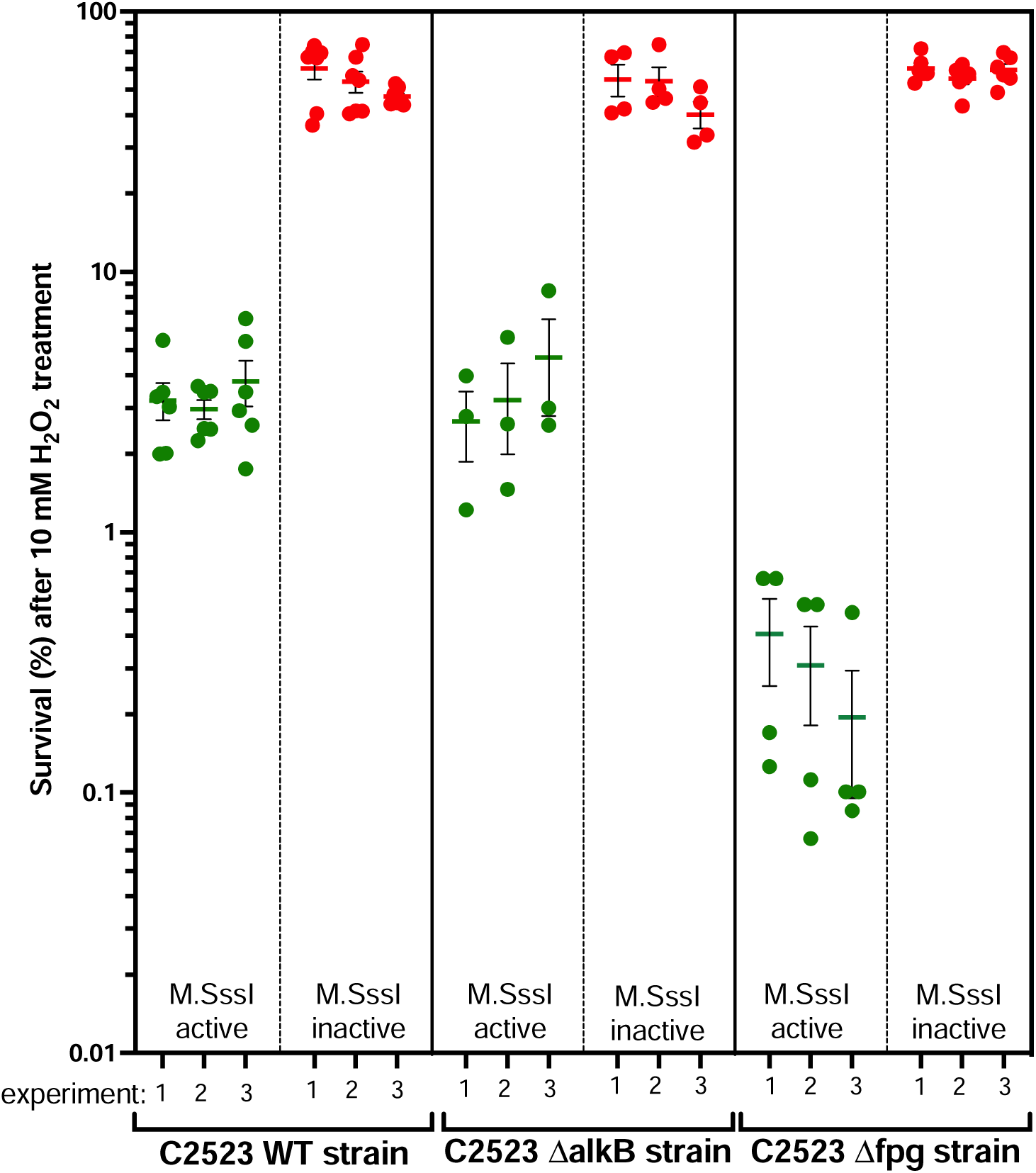
H_2_O_2_-induced sensitivity in strains carrying an active 5mC methyltransferase. WT strains and *alkB* and *fpg* mutants with M.SssI methyltransferase were treated with 10 mM H_2_O_2_. The percentage of survivals was calculated for each strain and normalized to the number of colonies growing under the same conditions without H_2_O_2_ treatment. Each experiment shows independent clones from the same transformation. Three separate transformations were performed for each plasmid.

### Increased ROS production in E. coli expressing DNMTs

We wondered why DNMT expression might induce sensitivity to oxidative stress. DNMT expression might lead to production of ROS directly. We sought to minimise secondary effects of long-term growth with DNMTs so designed a system whereby we could induce DNMT expression. We expressed M.SssI under the control of the pBAD promoter (methods). When grown in glucose, M.SssI remained silent, but was induced by adding Arabinose (Supplemental Figure 4). We measured ROS production using a commercially available fluorescent sensor (see methods). The ROS detection reagent in this system is DCFH-DA, a generalised ROS sensor that is not specific to any particular ROS molecule. We initiated expression of DNMT by adding arabinose and followed ROS accumulation over time under conditions in which baseline ROS accumulation was minimised (see methods and Supplemental Figure 5). As expected, H_2_O_2_ treatment led to an accumulation of ROS. H_2_O_2_-induced ROS accumulation was markedly enhanced by addition of DNMT(Figure 4). Moreover, DNMT addition alone led to increased ROS accumulation, even without exposure to H_2_O_2_ (Figure 4). Interestingly, the effect of H_2_O_2_ and ROS was greater than the linear combination of the two effects separately (linear model interaction P-value<0.05). Together these data strongly suggested that DNMT increased ROS levels, and that there was a synergistic effect with H_2_O_2_.

**Figure 4.**
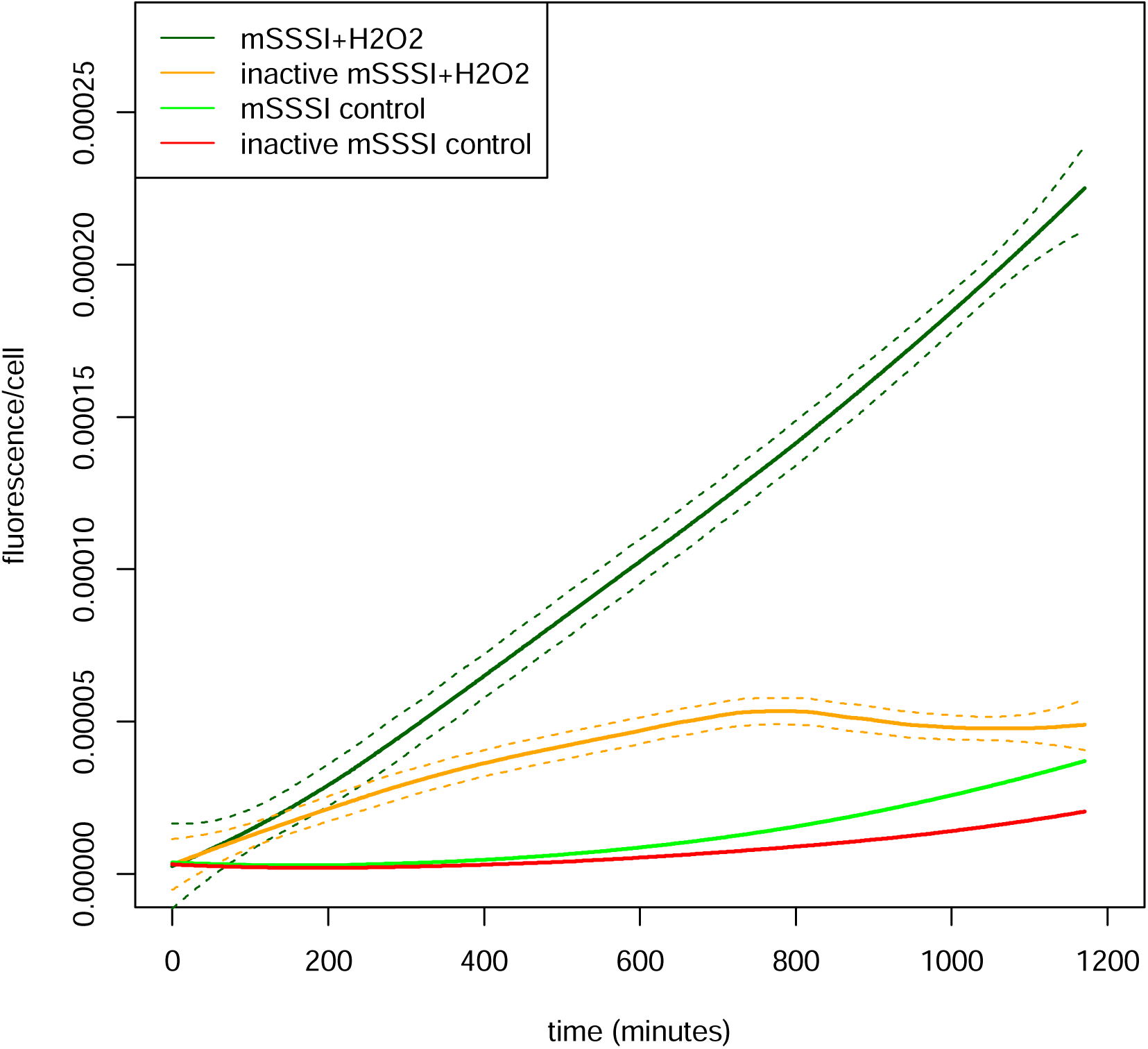
Reactive Oxygen Species production in the presence of M.SssI and H_2_O_2_. M.SssI was either induced with arabinose or kept repressed at the beginning of the experiment. Fluorescence per cell is plotted over 1200 minutes for each of the indicated conditions. The lines represent the loess fit from pooled data across two replicate experiments (two separate clones from a single transformation for each condition). The dashed lines show the standard error of the loess fit. Note that the error of the fit is too small to be visible as separate lines for conditions with very low reactive oxygen species.

The oxidative stress response relies on control of gene expression, in particular through the regulator oxyR which leads to transcriptional upregulation of a number of oxidative stress response genes including catalase which detoxifies H_2_O_2_ and reduces ROS production(Chiang & Schellhorn, 2012). Although 5mC is not recognised to have effects on transcription in *E. coli*, we considered the possibility that 5mC perturbed transcription of oxidative stress response genes, hence leading to failure to detoxify ROS. We therefore compared gene expression in the presence or absence of DNMT to test whether the transcriptional response to H_2_O_2_ treatment was impaired.

Induction of DNMT expression during logarithmic growth led to abundant transcriptional changes (Supplemental Figure 6A), suggesting that 5mC may have effects on gene expression. Despite our inducible system, some of these changes are likely to be secondary effects. For example, we observed upregulation of AlkB and its transcriptional regulator ada, although not aidB which is also regulated by ada, upon DNMT induction(Mielecki & Grzesiuk, 2014), which likely reflects production of alkylation damage in the form of 3mC by DNMT expression (Supplemental Figure 6A; Supplemental Table 4). The substantial transcriptional responses could potentially affect how individual cells respond to genotoxic stress and thus could be contributing to some of the excess sensitivity to MMS and H_2_O_2_ in cells expressing DNMTs. However, the induction of oxyR regulated genes such as catalase was unaffected by 5mC (Supplemental Figure 6B). Thus, the increased sensitivity to H_2_O_2_ is unlikely to be caused by failure of detoxification gene induction caused by DNMT expression.

### DNMT sensitizes cells to ROS through 5hmC and 5fC production

The synergistic effect of DNMT expression with deletion of Fpg (Figure 3) suggested that DNA damage might be implicated in the increased sensitivity to ROS exhibited by DNMT-expressing cells. One possibility is that oxidation of 5mC might result in damaged DNA bases, which would not occur in the absence of DNMT expression and thus might account for increased sensitivity to oxidative stress. Supporting this hypothesis we observed that exposure of DNMT-expressing cells to H_2_O_2_ led to production of the 5mC oxidation products 5hmC and 5fC as measured by mass spectroscopy (Figure 5). In eukaryotes enzymes of the TET family catalyse sequential oxidation of 5mC to 5hmC and 5fC(He et al., 2011; Ito et al., 2011; Tahiliani et al., 2009). *E. coli* does not have TET enzymes. However, ROS have been shown to act non-enzymatically on 5mC to convert it directly to either 5hmC or 5fC(Madugundu et al., 2014). 5hmC is not recognised as DNA damage, but 5fC is a substrate for base excision repair(Bochtler et al., 2017). Moreover, its presence in the DNA might indicate the presence of other 5mC oxidation products which our mass spectrometry did not measure, which could also be recognised by cells as DNA damage.

**Figure 5.**
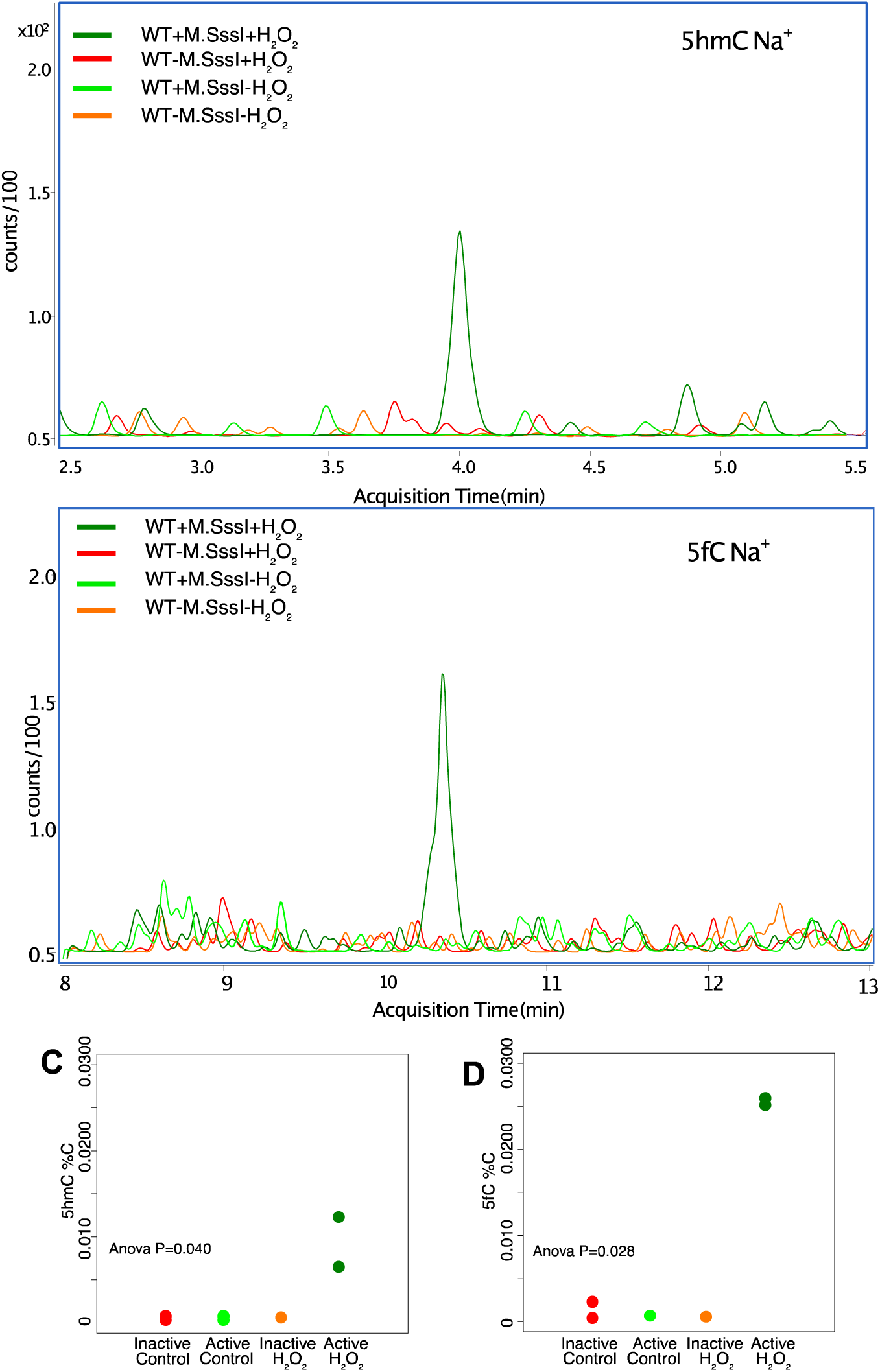
- Production of 5hmC and 5fC from upon H2O2 treatment in the presence of M.SssI. A and B show representative mass spectrometry traces for peaks identified as 5hmdC and 5fdC respectively. Quantitation is in figure C and D, where data points are the mean of two technical replicates and each data point is a separate clone from the transformation. Y-axis shows level of 5hmC or 5fC measured by mass spectrometry as a percentage of unmodified cytosine. P-value from a one way anova for effect of the experimental condition on the measurement is indicated on each plot.

We speculated that the production of 5fC and potentially other, downstream oxidation products of 5hmC or 5fC induced by H_2_O_2_ treatment might contribute to the excess sensitivity of DNMTs to H_2_O_2_ treatment. On the basis of this hypothesis, we predicted that artificially inducing 5hmC and 5fC production from 5mC would increase downstream oxidation products further and thus further increase DNMT sensitivity to ROS. To test this we expressed TET from *Naegleria gruberi*(Hashimoto et al., 2013) in *E. coli* under the control of an IPTG-inducible promoter, to induce sequential formation of 5hmC and 5fC from 5mC. Expression of NgTET in the presence of mSssI activity led to induction of both 5hmC and 5fC, confirming that NgTET was active and able to perform sequential oxidation of 5mC (Supplemental Figure 7). NgTET expression alone had a small but significant effect on sensitivity to H_2_O_2_ treatment, which may be due to the ability of NgTET to oxidise T to 5hmU(Pfaffeneder et al., 2014). However, when combined with DNMT expression there was a strong increase in sensitivity beyond what would be expected from either DNMT expression or NgTET expression alone (Interaction Tet*DNMT*IPTG p=1e-10 Anova; Figure 6A,B). Indeed, under the conditions of our assay there was essentially no growth at all when H_2_O_2_ was present-a synthetic lethal interaction (Figure 6A,B). We obtained similar results using a different TET enzyme, the phage-derived TET43 (Figure 6A). The production of 5hmC, 5fmC and further, downstream, oxidation products derived from 5mC may thus contribute to the excess sensitivity of cells expressing DNMTs to oxidative stress.

**Figure 6.**
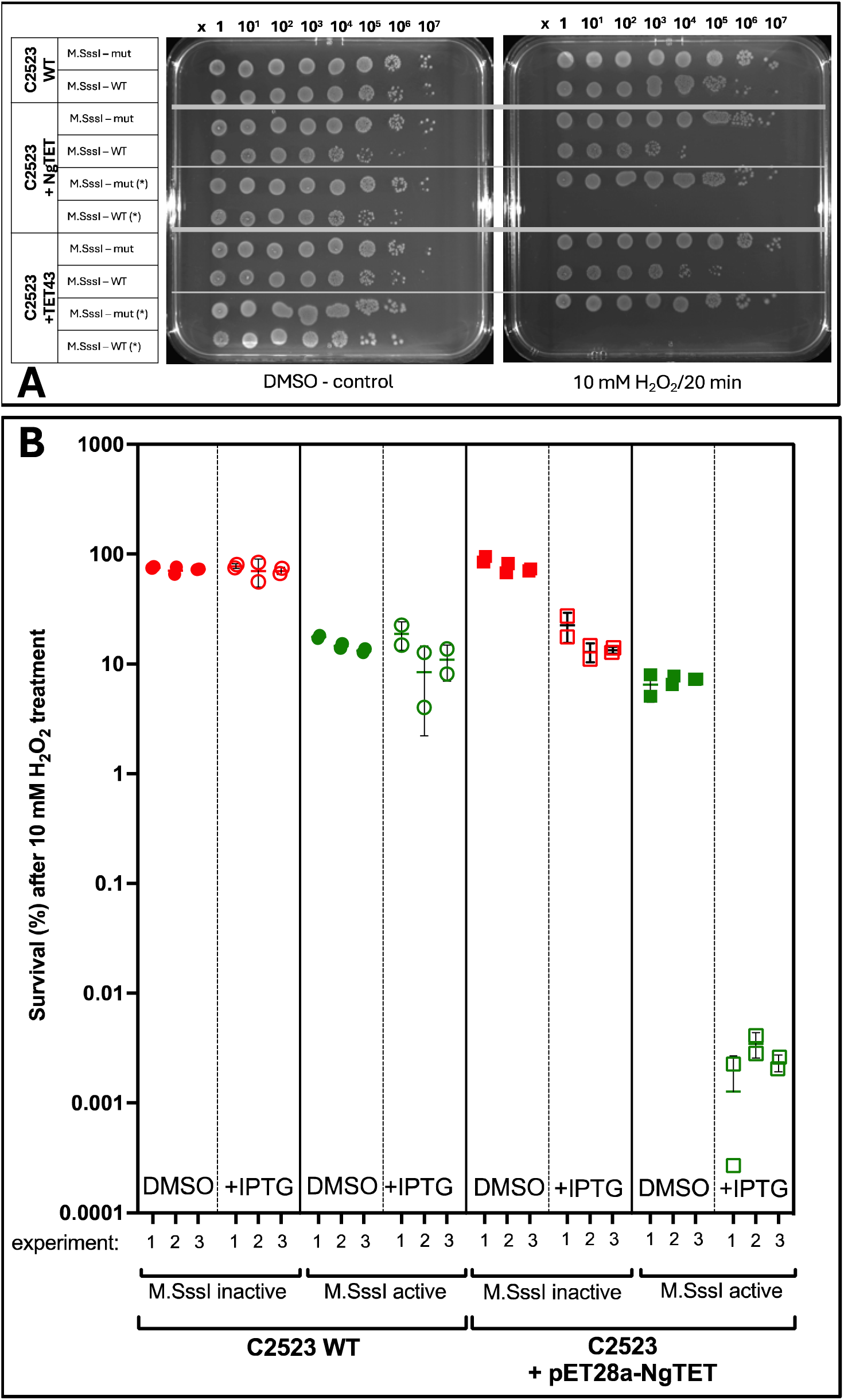
H_2_O_2_-induced sensitivity in strains carrying an active 5mC methyltransferase and expressing TET responsible for oxidative derivatives of 5mC. The WT strain and a strain carrying IPTG-inducible TETs (Negleria TET – NgTET or phage TET – TET43, more details in Materials section), which modifies 5mC to its oxidative derivatives that, in addition, also overexpressed inactive or active M.SssI methyltransferase, were treated with 10 mM H_2_O_2_. **A** – Drop plate test for *E. coli* cells without and after 10 mM H_2_O_2_ treatment (more technical details on Figure 3). **B** – The percentage of survivals from WT and NgTET cells was calculated for each strain and each condition and normalized to the number of colonies growing under the same conditions without H_2_O_2_ treatment. Each set of 3 points represents 3 independent clones from a single transformation and three separate transformations were performed.

## Discussion

In addition to its roles in regulating transcription epigenetically, cytosine-5 DNA methylation is increasingly recognised as a source of DNA damage. The heterologous DNMT expression system that we established in this manuscript enabled us to probe how the propensity of cytosine DNA methyltransferases to damage DNA affects sensitivity to exogenous genotoxic stress. Below we discuss how our findings indicate a novel connection between DNMTs and DNA damage through their impact on oxidative stress sensitivity. We also discuss the implications of our results for the evolution of DNMT activity and expression.

### Oxidative stress as a new source of DNA damage induction by DNMT expression

A major new finding of our paper is that DNMT expression sensitizes cells to oxidative stress. Importantly, we demonstrated that this does not proceed through the induction of DNA alkylation damage (3mC) by DNMTs because loss of AlkB activity did not exacerbate the sensitivity. What, then, might be the mechanism? One possibility is that DNMT activity itself generates reactive oxygen species, which increase the total cellular burden and making cells more susceptible to ROS production when H_2_O_2_ is added. However, the catalytic cycle of DNMT induction of 5mC is not known to involve radical generation ((Jeltsch, 2006)

An alternative explanation is that ROS causes toxicity through reaction of ROS with the major product of DNMTs (5mC) to generate DNA damage. We observed that H_2_O_2_ treatment increases the levels of 5hmC and 5fC in the DNA. Previously, free radical chemistry has been demonstrated to lead to formation of these oxidation products(Madugundu et al., 2014), thus we propose that the same process occurs in our system when H_2_O_2_ is added.. The relatively higher levels of 5fC rather than 5hmC that we observed is consistent with this suggestion as OH· radicals were shown to produce 5fC directly from 5mC(Madugundu et al., 2014), in contrast to the enzymatic oxidation which proceeds sequentially(Ito et al., 2011). In mammals 5fC is a lesion in the DNA which is excised by base excision repair, most prominently by the activity of TDG aided by NEIL1 and NEIL2(Bochtler et al., 2017). Whilst no direct counterpart of TDG exists in *E. coli*, mismatched-uracil glycosylase (MUG) is structurally similar. Moreover, NEIL1 and NEIL2 are related to Fpg in *E. coli*(Hwang et al., 2025; Jacobs & Schär, 2012). We provided further evidence for this model by accelerating 5fC formation through co-expression of TET with DNMT, which resulted in synthetic lethality upon exposure to H_2_O_2_. Importantly, although our mass spectrometry investigations were limited to 5hmC and 5fC their presence likely indicates that other oxidation products of 5mC would be created which would also be substrates for glycosylases(Slyvka et al., 2017). This explanation would suggest that the excess sensitivity to oxidative stress caused by DNMT expression is due to 5mC being oxidised to form modified nucleotides that may be recognised and excised as DNA damage (Figure 7).

**Figure 7.**
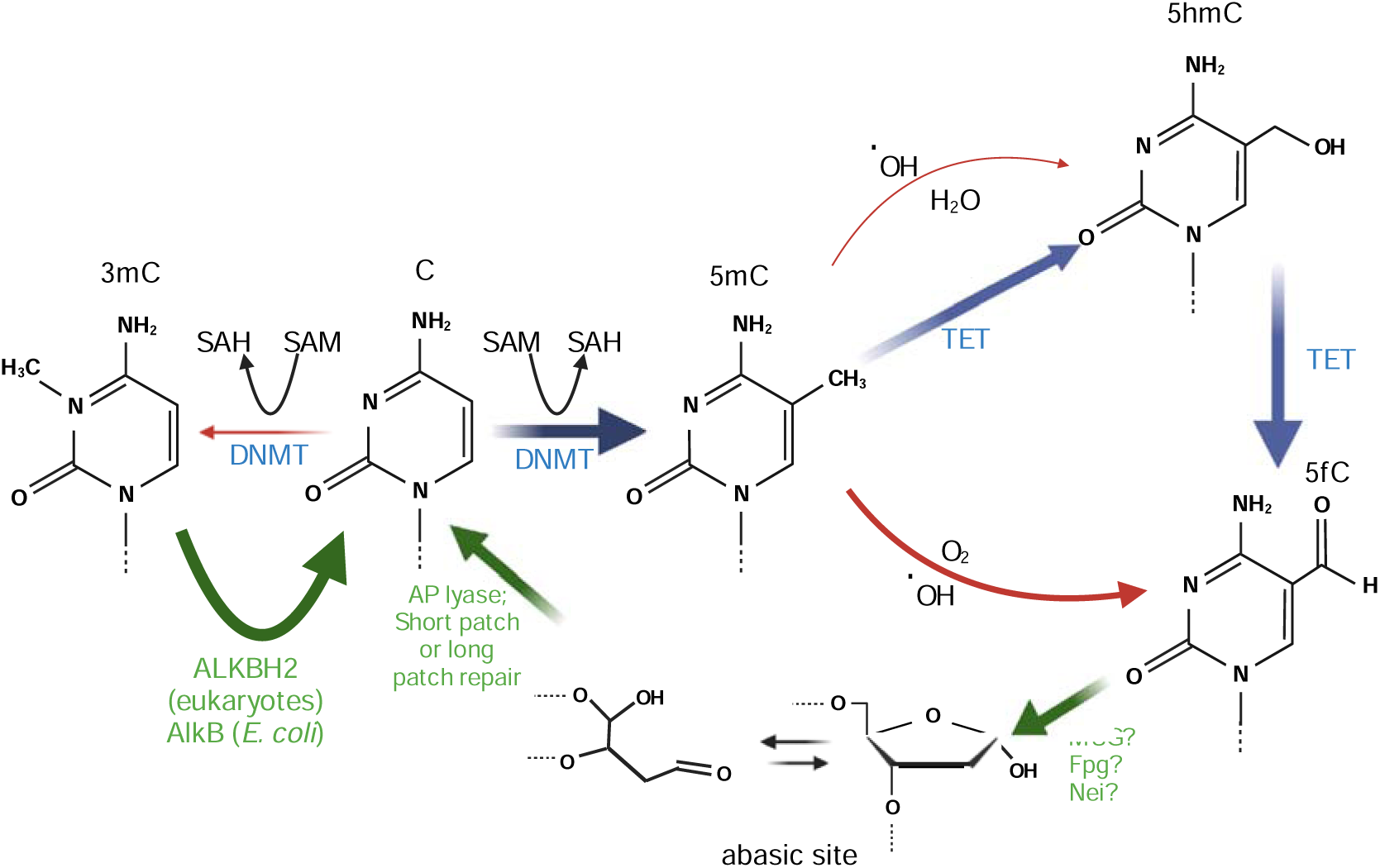
- Model: sources of DNA damage due to DNMT expression. Repair pathways are shown in green, enzymatic pathways in blue and DNA damage mechanisms in red.

We suggest that this mechanism may also explain why we observed that ROS levels increase when cells expressing DNMTs are exposed to H_2_O_2_. Reaction of 5mC with ROS proceeds via a radical cytosine derivative, with alternative pathways potentially generating other free radicals via reactions with oxygen or water in addition to the formation of 5mC and 5hmC(Madugundu et al., 2014). As a result, it is possible that the presence of 5mC itself could be sufficient to result in increased levels of ROS. Such a scenario would directly link the increased ROS levels we observed in DNMT-expressing cells to the predominant product of DNMT activity. We believe this is a plausible mechanism to explain both increased ROS and increased sensitivity to oxidative stress when DNMT is expressed. However, other explanations are possible, and it is notable that DNA damaging agents such as MMS can lead to ROS generation(Rowe et al., 2008). A more detailed chemical and kinetic study of the ROS formation in DNMT-expressing cells would be needed to resolve these questions.

The observation that TET overexpression sensitizes cells expressing DNMTs to oxidative stress strongly suggests that the site of DNA damage is the modified cytosine itself. However, we do not currently have definitive evidence supporting this. As mentioned in the results section, the presence of unrepaired 3mC may lead to increased levels of ssDNA; it is also possible that 5mC itself may increase ssDNA levels. Loss of alkB would be expected to increase the amount of ssDNA. Thus DNA damage surrounding modification sites, but not specifically localised to it, might be the cause of the increased sensitivity. These two different models make different predictions. If modified cytosines are the source of the damage, mutations arising would be predominantly located at CG dinucleotides. Alternatively, ssDNA exposure would result in distributed mutations that would not necessarily be located at CG sites. The highly biased spectrum of mutations that can be screened through the Rif resistance assay does not allow us to address this currently. However, future experiments to create mutation accumulation lines could allow us to address the question systematically on a genome-wide level.

Although we favour the interpretation that oxidative stress is a direct consequence of DNMT activity in *E. coli*, it is important to note that we did observe an unexpectedly large number of gene expression changes due to mSSSI expression (see supplemental Table 4), Although no salient downregulation of ROS processing machinery were observer, we cannot exclude that altered transcription could contribute to increased ROS sensitivity. The source of the extensive gene expression changes, including the possibility that extensive cytosine methylation might directly perturb *E. coli* transcription, is worth investigating further.

### Implications of the effect of DNMTs on sensitivity to genotoxic stress for the evolution of DNMTs

The key enzymes that introduce and maintain cytosine DNA methylation in animals are found across the eukaryotic tree indicating an ancient origin at the dawn of eukaryotic cells(Ponger & Li, 2005). However, the evolution of DNA methyltransferase pathways across eukaryotes is strikingly rapid, with many different lineages having lost DNMTs completely(de Mendoza et al., 2020). To what extent might the propensity of DNMTs to damage DNA be responsible for the frequent losses of DNMTs? Due to the coevolution of DNMTs with the alkylation repair enzyme ALKBH2, we speculated that DNMT’s off-target activity may pose a threat to genome stability which imposes a fitness cost, even in the presence of the ALKBH2 repair pathway(Sarkies, 2022a). A key prediction of this model is that DNMT expression would be disadvantageous to cells in the absence of ALKBH2 activity. Our *E. coli* model of DNMT expression enabled us to test this and indeed we found that deletion of AlkB leads to a large increase in sensitivity to the alkylating agent MMS when combined with DNMT expression. Thus the propensity of DNMTs to alkylate DNA could lead to a fitness cost at least in the presence of environmental sources of alkylation damage. It is notable that there is also a small decrease in fitness due to combined expression of DNMT and deletion of AlkB, even without MMS treatment. Nevertheless, for this to make loss of DNMT expression favourable the benefit of DNMT expression in terms of other functions such as gene expression would have to be quite small.

High concentrations of alkylating agents such as MMS are not a common component of ecologically relevant niches. However, oxidative stress is much more commonly experienced by organisms(Samet & Wages, 2018). Our observation that DNMT expression renders *E. coli* hypersensitive to oxidative stress may therefore provide a plausible reason why organisms with DNMTs might be at a selective disadvantage in certain environments. Switching into an environment with higher levels of oxidative stress might therefore result in the cost of DNMT expression outweighing its benefit leading, over evolutionary time, to its loss. It would be very interesting to establish the environments of species with and without DNMTs to test whether exposure to oxidative stress might be associated with high rates of DNMT loss.

## Materials and methods

### Bacterial strains

*Escherichia coli* B wild-type (WT) strain C2523 which does not cleave methylated DNA (*mcrA^—^*, *mcrBC^—^*, *mrr^—^*, *dcm^—^*, *dam^+^*) with the genotype *fhuA2 [lon] ompT gal sulA11 R(mcr-73::miniTn10--Tet)2 [dcm] R(zgb-210::Tn10--Tet) endA1 Δ(mcrC-mrr)114::IS10* (New Engalad Bioloabs, NEB) was used in this study. Single deletions of *alkB* gene *(ΔalkB::kan^R^*) and *fpg* gene *(Δfpg::kan^R^*) were carried out by P1-mediated transduction using the corresponding knockout strains from the Keio collection as donors(Baba et al., 2006). In this system the targeted gene is replaced with the kanamycin gene flanked by the FRT sequence for subsequent removal of kanamycin cassette. To remove the kanamycin cassette, pCP20, an ampicillin and chloramphenicol resistant plasmid with temperature-sensitive replication and flippase recombinase synthesis(Cherepanov & Wackernagel, 1995) was transformed into the *ΔalkB::kan^R^*mutant and selected at 30°C, after which pCP20 was purified at 43°C. Removal of knockout genes and the antibiotic resistance gene were confirmed by PCR using and by culturing the strains in antibiotic containing LB media.

To control DNA methyltransferase expression from arabinose inducible promotor *E. coli* TOP10 K12 (NEB) strain with the genotype *F-mcrA (mrr-hsdRMS-mcrBC) 80lacZ M15 lacX74 recA1 ara 139 (ara-leu)7697 galU galK rpsL (StrR) endA1 nupG* was used. This strain also does not cleave 5mC methylated DNA.

### Plasmids

The bacterial CpG DNA 5-methylcytosine (5mC) methyltransferase from *Spiroplasma sp*. (M.SssI) (UNIOROT accession number P15840) cloned into plasmid pAIT2, a low copy number plasmid with kanamycin (Kan) and chloramphenicol (Chl) resistance was used in this study. M.SssI is expressed under lacZ promoter. The sequence of M.SssI ORF is 1158 bps long and has 386 amino acids(Renbaum et al., 1990). pAIT2 with truncated M.SssI (M.SssI-inactive) was created by removing the first 570 bps of M.SssI, which includes the SAM binding domain of the protein. The full length of M.SssI coding sequence was re-cloned into pBAD plasmid with ampicillin (Amp) resistance to control DNA methyltransferase expression from arabinose (P_BAD_) promoter by addition of arabinose (ara) to the media(Guzman et al., 1995). Plasmid with bacterial CpG DNA 5-methylcytosine (5mC) methyltransferase from *Mycoplasma penetrans* (M.MpeI) was kindly provided by Matthias Bochtler (IIMCB, Warsaw, Poland). Briefly, the 1185 bp sequence encoding 395 aa long M.MpeI protein was cloned into pET-28a (Novagen, Km) plasmid. This M.MpeI variant differed from the UNIPROT protein (accession Q8EVR5) by the Q68R, K71R, S295P point mutations without affecting an enzymatic properties(Wojciechowski et al., 2013).

The degree of CpG methylation from pAIT2-M.SssI-active, pAIT2-M.SssI-inactive, pBAD-M.SssI-active and pET28a-M.MpeI-active plasmids were assessed by restriction endonuclease digestion with the methylation sensitive enzyme, HpaII (CCGG target sequence), or the methylation insensitive isoschizomeric endonuclease, MspI and subsequent agarose gel electrophoresis monitoring visualized by GelDoc system (BioRad). The heterolobosean amoeboflagellate *Naegleria gruberi* ten-eleven translocation (TET) methylcytosine dioxygenase (NgTET) cloned into IPTG-inducible pJS119K plasmid (Km) and the phage-derived TET43(Burke et al., 2021) were kind gift from Lana Saleh (New England Biolabs, Massachusetts, USA).

### Bacterial media and culture conditions

All strains were cultured in Luria-Bertani (LB) medium (1% NaCl, 0.5% yeast extract, 1% tryptone) with or without appropriate antibiotics (Kanamycin (Kan) 50 μg/ml, Ampicillin (Amp) 100 μg/ml, Chloramphenicol (Chl) 30 μg/ml) at 37°C with 220 rpm. Similarly, the bacteria were plated in LB plates (1.6% agar) with or without the aforementioned concentrations of antibiotics and incubated at 37°C overnight.

In order to tightly control expression of M.SssI methyltransferase under arabinose promoter P_BAD_ the strain TOP10 with pBAD-M.SssI-WT plasmid was cultured in LB supplemented with 0.5% glucose (to hold the expression) or 0.5% L-arabinose (to induce protein production) additionally supplemented with Amp. Briefly, a strain with inducible DNA methyltransferase was cultured overnight in LB medium supplemented with 0.5% glucose to maintain promoter suppression. The bacteria were then diluted 1:100 and a new culture was started under the same conditions to an OD600 of about 0.3. The bacteria were then centrifuged and washed, and the medium was changed to LB with 0.5% arabinose to induce protein expression from the P_BAD_ promoter.

### Treatment with DNA damaging agents

#### MMS acute treatment

Sensitivity to alkylation DNA damage was tested for C2523 WT and *ΔalkB* strains expressing either low (pAIT2) copy number plasmids containing M.SssI or its inactive version (M.SssI-tc) or high copy number plasmid (pET28a) containing active M.MpeI. All experiments were repeated at least 3 times, each time with 3 repeats of each condition, always including WT and mutant parent strains. Bacteria were cultured in LB medium with appropriate antibiotics to OD600 0.6 to 0.8. One half of the culture served as untreated control and the other was treated with 20 mM methylmethane sulphonate (MMS, Sigma) for 20 min, during which both sets of the bacterial culture continued to grow at 37°C with 220 rpm. After treatment, cultures were directly and quickly diluted in PBS to allow visualization of individual colonies and placed on LB agar plates supplemented with the appropriate antibiotic. After overnight incubation at 37°C, the colonies were counted and the colony forming unit (CFU/ml) was calculated. To normalize the effect of MMS, the percentage of survival was calculated by taking the number of bacteria growing under control conditions as 100% and calculating the percentage surviving after mutagen treatment.

#### MMS titration

To test the sensitivity to MMS over a range of concentrations, C2523 WT and *Δ*alkB strains expressing active M.SssI DNA methyltransferase (M.SssI-WT) or its inactive version (M.SssI-mut) were exposed to different times and doses of MMS. All tests were repeated 3 times, each time with 2 replicates of each condition, always including the WT and mutant parental strains. Bacteria were grown in LB medium with the appropriate antibiotics to an OD600 of 0.6. One aliquot of untreated cells was used as a control, and the other was exposed to different doses of MMS (0, 5, 10, 20, 30, and 60 mM) for 20 min or different times of action of 20 mM MMS (0, 5, 10, 15, 20, and 30 min). All sets of bacterial cultures were continued to grow at 37°C at 220 rpm. At each time, the cultures were directly and rapidly diluted in PBS, and a small aliquot was used for the drop plate test to visualize the effect (described belove), and the rest were plated onto LB agar plates supplemented with the appropriate antibiotic. After overnight incubation at 37°C, colonies were counted to obtain the percentage of survival after mutagen treatment, and the remaining plates were recorded as images to show the effect of MMS titration dose and time dependence.

#### Cisplatin and H_2_O_2_ treatment

Cisplatin and H_2_O_2_ were carried out following the protocol for MMS acute treatment. Cisplatin (Merck) was dissolved in DMF (Sigma) and the cells were treated with 100 μM cisplatin for 1h. 30% solution of H_2_O_2_ (Sigma) was used to treated bacteria with final concentration of 10 mM H_2_O_2_ for 20 min.

Statistical analysis was performed in R. A two way anova was used to test the effect of M.SssI expression on sensitivity to DNA damaging agents and a three way anova used to incorporate the effect of specific mutations on the effect of M.SssI. Presence of a synthetic interaction was determined by significance of the relevant interaction term such that the combined effect of the mutant and the M.SssI on sensitivity was greater than additive i.e. greater than the sum of the mutant and M.SssI alone. The code used took the form: Model<-aov(growth∼drug*M.SssI*mutant) where %growth is the number of colonies normalised to the WT control from the same experiment.

### Drop plate test for quick enumeration

The drop plate method after mutagen treatment was perform for quick enumeration of *E. coli* cells.

WT strains and *alkB* and *fpg* mutants with M.SssI inactive and active methyltransferases were cultured to reach OD600 of 0.4 and then treated with: DMSO (as control), 20 mM MMS, 10 mM H_2_O_2_ or 100 μM cisPt, next the series of dilutions were made, and the drops (5 μl) were plated on LB (Chl/Kan) square plates. The dose of spots from left to right corresponded to 1, 2, 3, 4, 5, 6, 7 log10 dilutions of *E. coli*.

### Mutation rate assessment via Rifampicin resistance assay

Spontaneous mutation frequency was tested on LB agar plates containing 100 μg/ml rifampicin (Rif-plate). After fresh plasmids (encoding active and inactive M.SssI) transformation into C2523 WT and *alkB* mutant cells single colonies were picked and used to inoculate independent 3 ml LB cultures supplemented with kanamycin and chloramphenicol. Cultures were grown with agitation overnight at 37 °C and appropriate dilutions were spread on LB plates (Chl/Kan) to determine the total cell count and on LB agar plates containing 100 μg/ml rifampicin to determine the number of rifampicin-resistant mutants (RifR). LB (Chl/Kan) plates were incubated overnight, and Rif-plates were incubated at 37 °C. for 48 h. Mutation frequencies were calculated by dividing the number of colonies formed by mutants by the total number of colonies. For spontaneous mutation frequency determinations, each experiment was repeated 3 times with 3 biological replicates each time (a total of 9 biological replicates). A drop test was also used to visualise qualitatively the dependence of the spontaneous mutation rate on active M.SssI in an *alkB*-deficient background strain. The series of dilution from overnight cultures were made, and the drops (5 ml) were plated on LB (Chl/Kan) and LB (Rif) square plates. The dose of spots from left to right corresponded to 1, 2, 3, 4, 5, 6, 7 log_10_ dilutions of *E. coli*.

### Reactive Oxygen Species (ROS) production monitoring

WT Top10 cells were made competent and transformed with the pBAD M.SssI construct or dH_2_O. The transformed bacteria were plated on LB High Salt Agar + 0.8% Glucose (+100µg/mL Carbenicillin for pBAD M.SssI transformants) and incubated overnight at 37⁰C. Individual colonies from Top10 and Top10 + pBAD M.SssI plates were then inoculated overnight in LB High Salt media +0.5% Glucose (+100µg/mL Carbenicillin for pBAD M.SssI transformants). The next day, cultures were diluted 25X and grown to OD600 = 0.4. 3ml of the original overnight culture was taken for plasmid DNA extraction and methylation analysis. Cultures were then split into 3×6ml, centrifuged at 4700 rpm for 3 mins and resuspend in LB High Salt media + 0.5% Arabinose, 0.5% Arabinose + 0.15% Glucose or 0.15% Glucose respectively. Cultures were grown to OD600 = 0.6 and split into 2×3ml. Cultures were treated with 10mM or 0mM H2O2 then incubated at 37⁰C, 200rpm for 30 mins. 3ml was then taken for plasmid DNA extraction and methylation analysis. In a 96 well plate, 2 x 100µL was plated for each condition. 2µL 50X ROS Dye from the Total Reactive Oxygen Species (ROS) Assay Kit (520nm, Cat. No. 88-5930; Thermofisher) was added to each well. The plate reader was heated to 37⁰C and fluorescence measured by excitation at 475nm and emission at 500-550nm. Absorbance was then measured by OD600. The plate was then shaken for 10 mins at 200 cycles/min (orbital, 2mm shaking diameter). This loop was repeated 100 times.

### Genomic DNA isolation and Mass Spectrometry analysis of DNA epigenetic modifications

C2523 WT cells expressing M.SssI methyltransferase or its inactive version (M.SssI-tc) as well as M.MpeI metyltransferase, both H_2_O_2_ treated and untreated, were cultured overnight in 10ml of LB at 37C with shaking. Cells were lysed and proteins simultaneously denatured in the appropriate lysis buffer according to the commercial kit DNeasy Blood & Tissue Kit (Qiagen, cat. no. 69504). Proteinase K then was added and after incubation in 55 °C for 30 min, lysates were loaded onto the mini columns and centrifuged. After several washing steps according to the protocol, pure, high-molecular-weight DNA was eluted and precipitated with isopropanol. Then, precipitates were diluted in water and 1 μg of pure DNA was checked for its quality and quantity on Genomic DNA ScreenTape (Agilent, cat. no. 5067-5365) using TapeStation 4150 (Agilent).

### DNA nucleoside analysis by mass spectrometry (HPLC–QQQ)

1500ng of DNA was treated with 5U DNA Degradase Plus™ (Zymo Research, E2020) in 120μl 1X DNA Degradase Plus™ Reaction Buffer. The mixture was incubated at 37°C for four hours and then lyophilised by SpeedVac to a volume of 3μl. 14.5μl of bufferA (10 mM ammonium acetate, pH 6) was added and the mixture was then filtered through a 96-well filter microplate with a 0.2µm polyethersulfone membrane via centrifugation at 3200g for 35 minutes at 4°C. Finally, a further 4μl of bufferA was added to the approximately 10μl of filtrate following centrifugation.

For the analysis by HPLC–QQQ mass spectrometry, a 1290 Infinity UHPLC was fitted with a Zorbax Eclipse plus C18 column, (1.8μm, 2.1mm 15mm; Agilent) and coupled to a 6495a Triple Quadrupole mass spectrometer (Agilent Technologies) equipped with a Jetstream ESI-AJS source. The data were acquired in dMRM mode using positive electrospray ionisation (ESI1). The AJS ESI settings were as follows: drying gas temperature 230°C, the drying gas flow 14 lmin, nebulizer 40 psi, sheath gas temperature 400°C, sheath gas flow 11 lmin, Vcap 2,000 V and nozzle voltage 0 V. The iFunnel parameters were as follows: high pressure RF 110 V, low pressure RF 80 V. The fragmentor of the QQQ mass spectrometer was set to 380 V and the delta EMV set to +200. The gradient used to elute the nucleosides started by a 5-min isocratic gradient composed with 100% bufferA (10 mM ammonium acetate, pH 6) and 0% buffer B (100% methanol – 106035, Merck) with a flow rate of 0.400 ml min 1and was followed by the subsequent steps: 8–9 min, 94.4 % A; 16 min 86.3 % A; 17– 21 min 0 % A; 24.3–25 min 100 % A. The gradient was followed by a 3-min post time to re-equilibrate the column. The raw mass spectrometry data was analysed using the MassHunter Quantitative Analysis Software package (Agilent Technologies, version 10.0). The transitions and retention times used for the characterization of nucleosides and their adducts are summarized in1[table ref]. For the identification of compounds, raw mass spectrometry data was processed using the dMRM extraction function in the MassHunter software. For each nucleoside, precursor ions corresponding to the M-H ^+^and M-Na ^+^species were extracted, and the signal observed was quantified against a standard curve. The chromatograms were generated using the MassHunter Qualitative Analysis Software package (Agilent Technologies, version 10.0).

### Induction of DNMT activity and preparation of cells for transcriptomics

Bacterial TOP10 cells with pBAD-M.SssI-WT plasmid or empty pBAD plasmid were cultured to stationary phase in 5 ml LB supplemented with 0.1% glucose to maintain promoter suppression and further supplemented with Amp. The cells were then diluted 1:100 and re-cultured to logarithmic phase (OD600 approximately 0.3) in 20 ml LB supplemented with 1% glucose and Amp, centrifuged, washed with glucose and divided in half. One half (10 ml) was cultured in LB with 1% glucose and Amp to maintain suppression of the P_BAD_ promoter, and the other 10 ml was cultured in LB with 0.1% arabinose and Amp to induce expression of DNA methyltransferase 5mC from the P_BAD_ promoter. During the 2 hours of growth, cells growing on glucose were diluted several times after reaching an OD600 of about 1 to 0.5, while cells growing on arabinose and producing M.SssI were unable to double their amount due to the production of a protein that was toxic to them and remained at the OD600 about 0.3). Nevertheless, the cultures were again divided in half, one half (5 ml) was left as a control and the other half (5 ml) was treated with 10 mM H_2_O_2_ for 20 minutes. After this time, all cultures were centrifuged, glucose or arabinose and H_2_O_2_ were washed off, and cells were resuspended in 250 μ of PBS, and next used for RNA isolation to RNA sequencing.

### RNA isolation

RNA was isolated using TRIzol purification technique. Briefly, the cell pellet was resuspended in 250 μl of PBS, then 1 ml of TRIzol reagent was added per sample. After 5 min of incubation at room temperature (RT) 0.2 ml of chloroform was added followed by 15 s variously shaking and incubation at RT for 3 min. Next, sample was centrifuged in microfuge on maximal speed for 15 min at 4 °C and the aqueous phase (top) of the sample was place into a new tube followed by 0.5 ml addition of 100 % isopropanol for RNA precipitation. After 30 min incubation in RT sample was pelleted by centrifugation at maximal speed for 15 min at 4 °C. The pellet was washed twice with 1 mL of 75% ethanol. Air dried the RNA pellet was resuspended in in RNase-free water (100 μL). Next, RNA was treated with 1 unit of DNaseI (DNAse I recombinant, RNase-free, ROCHE) per sample, incubated at 30 °C for 20 min and transferred to ice. DNaseI was removed from sample by mixing the sample with 350 μl of RLT buffer (QIAGEN RNeasy Mini kit) followed by addition of 350 μl ethanol (96 %) without centrifugation. 700 μl of sample was transferred into RNeasy Mini spin column (QIAGEN RNeasy Mini kit, cat. no. 74104), centrifuged for 15 s at 8000 x g. RNA on mini-column was twice washed with 500 μl of RPE buffer (QIAGEN RNeasy Mini kit). RNA was removed from the column by 30 μl of RNase free water. Quality and quantity of RNA was checked on RNA ScreenTape (Agilent, cat. no. 5067-5576) using TapeStation 4150 (Agilent). 1 μg of RNA was sent for RNA sequencing (Novogene).

### Transcriptomic data analysis

RNA sequencing reads were aligned to the *E. coli* genome using bowtie2 using the preset – very_fast. SAM files were processed to BAM files using samtools (Li) and bam files processed to bed files using bedtools. Counts for each annotated *E. coli* gene were calculated by intersecting the *E. coli* gene coordinate gff file with the bed file using bedtools. Subsequently, data analysis was performed in R, using DESeq2 to normalize the count data across samples and calculate log2 fold change for the different comparisons. Scripts for the R analysis are provided on the Sarkieslab github page: https://github.com/SarkiesLab/EcoliM.SssI_geneexpression

## Acknowledgements

We thank Dr Lana Saleh and Professor Matthias Bochtler for sharing plasmids used in this experiment. We thank Dr Ben Berks for help with the pBAD expression system and Dr Stefan Uphoff for sharing clones from the Keio library for knockout generation.

---

**Supplemental Figure 1.**
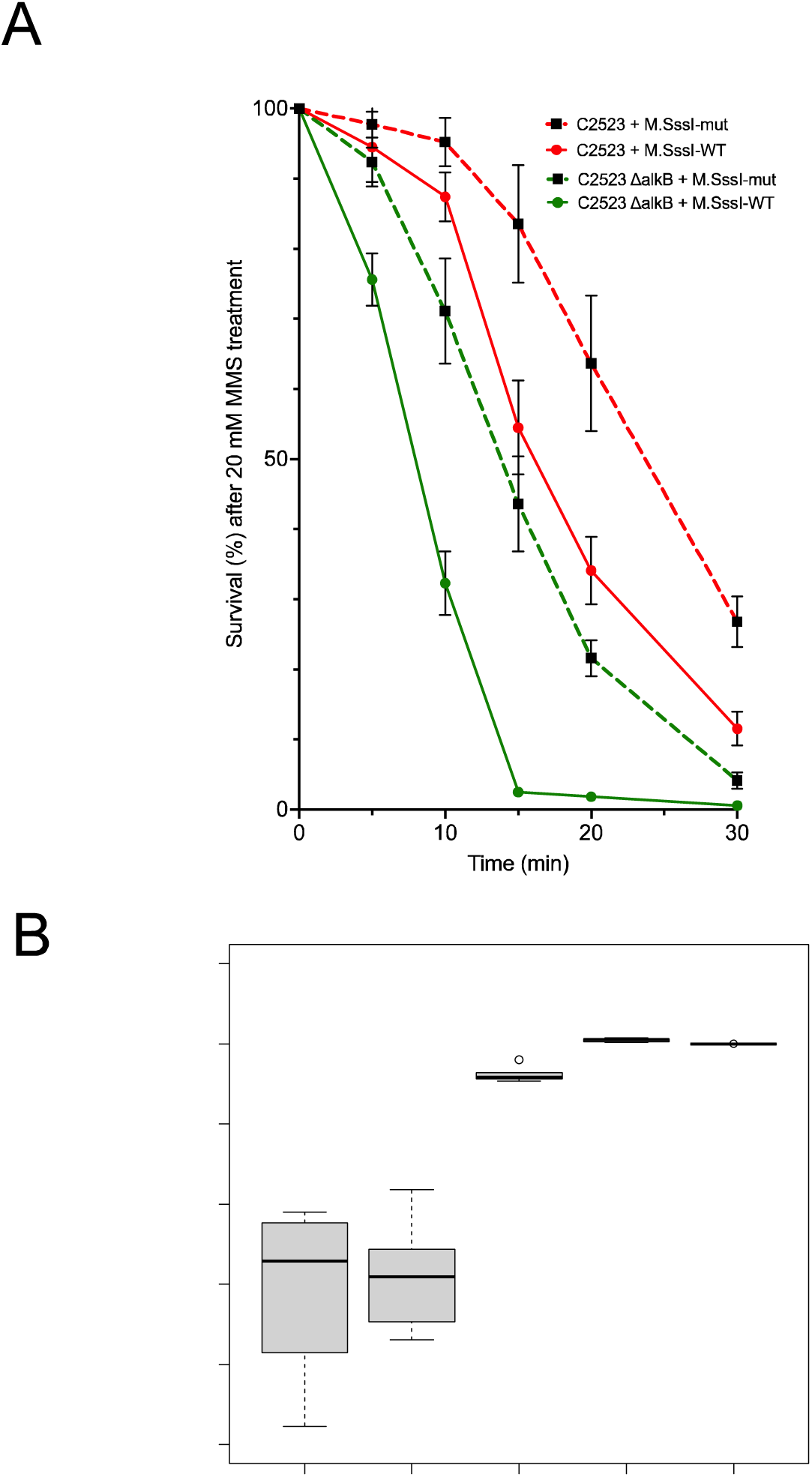
- MMS titration. A shows dose response curve for mSssI expressing E. coli cells either in a WT or *alkB* mutant background. B shows the fractional deviation from an additive interaction, demonstrating that at low doses a greater than additive effect on survival between expression of mSSSI and alkB mutation is observed. P values are from a two-tailed anova reporting an interaction term between mSssI expression and alkB mutation.

**Supplemental Figure 2.**
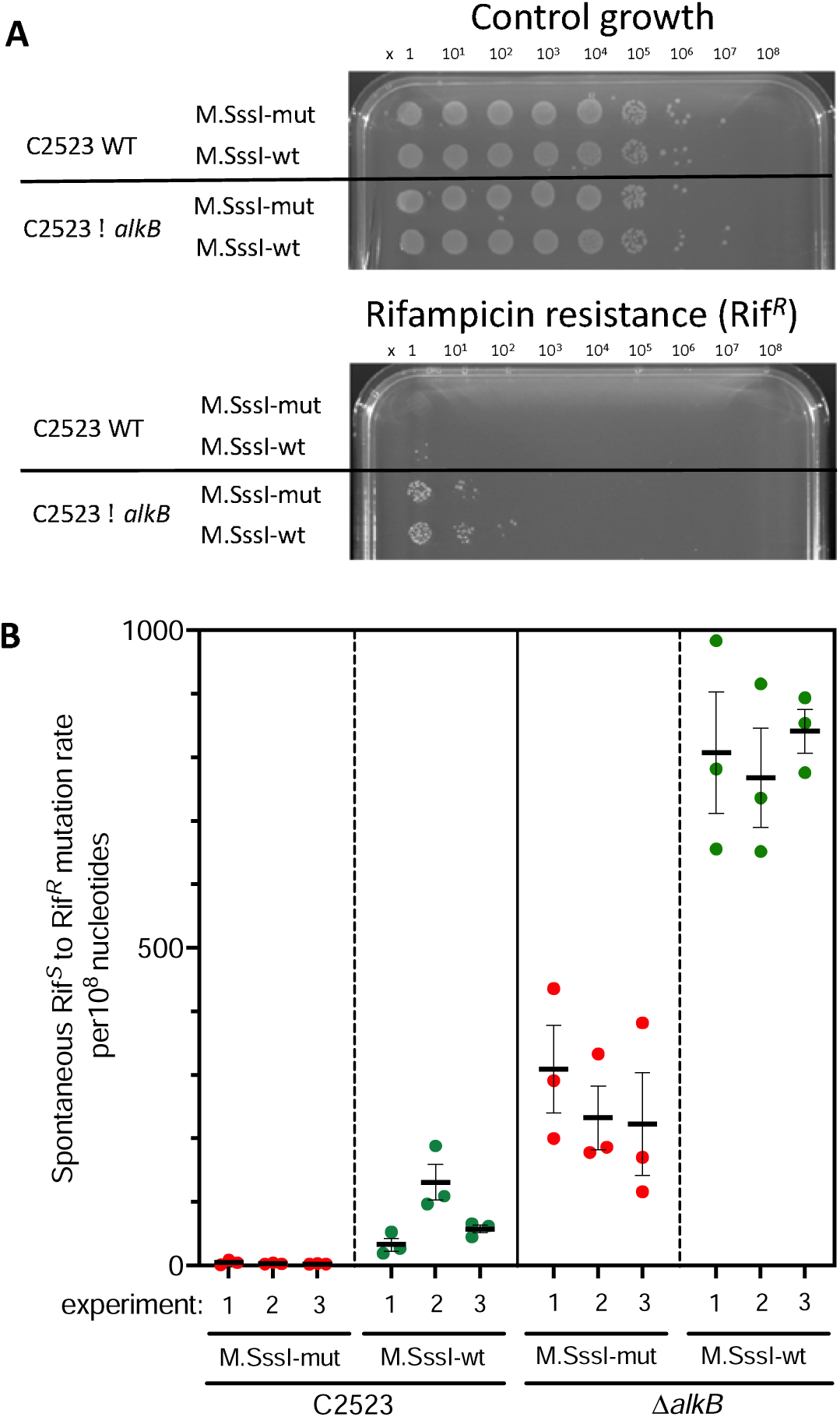
- Mutation rate in the presence or absence of mSssI in either WT or *alkB* mutant E. coli cells. Spontaneous mutation rate was measured using a rifampicin resistance assay (see methods).

**Supplemental Figure 3.**
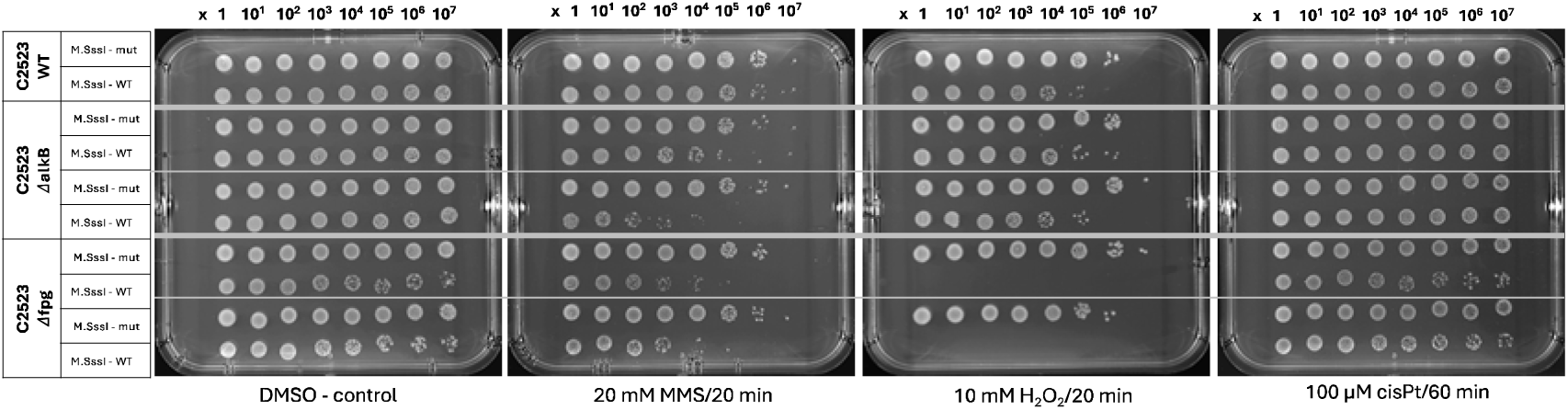
- Drop test screening sensitivity of M.SssI expressing cells to various DNA damaging agents. WT, *alkB* mutants and *fpg* mutants are shown.

**Supplemental Figure 4.**
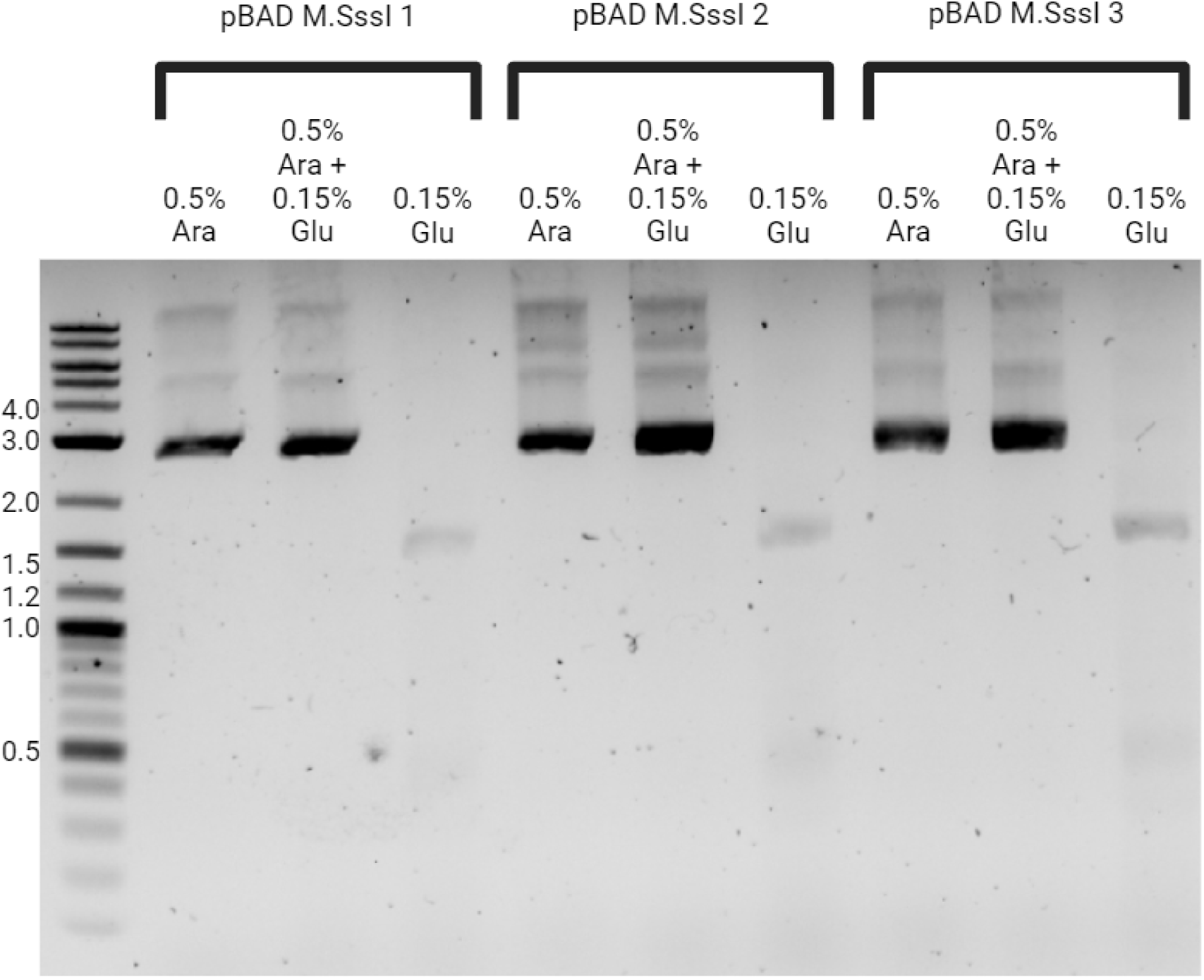
- Induction of M.SssI expression under control of pBAD promoter by arabinose. The plasmid was digested with methyl-sensitive restriction enzyme HpaII. The presence of the upper band indicates undigested plasmid. Arabinose maintains methyltransferase expression even in the presence of 0.15% glucose, whereas removal of arabinose leads to repression of the methyltransferase. Three independent clones from a single transformation are shown.

**Supplemental Figure 5.**
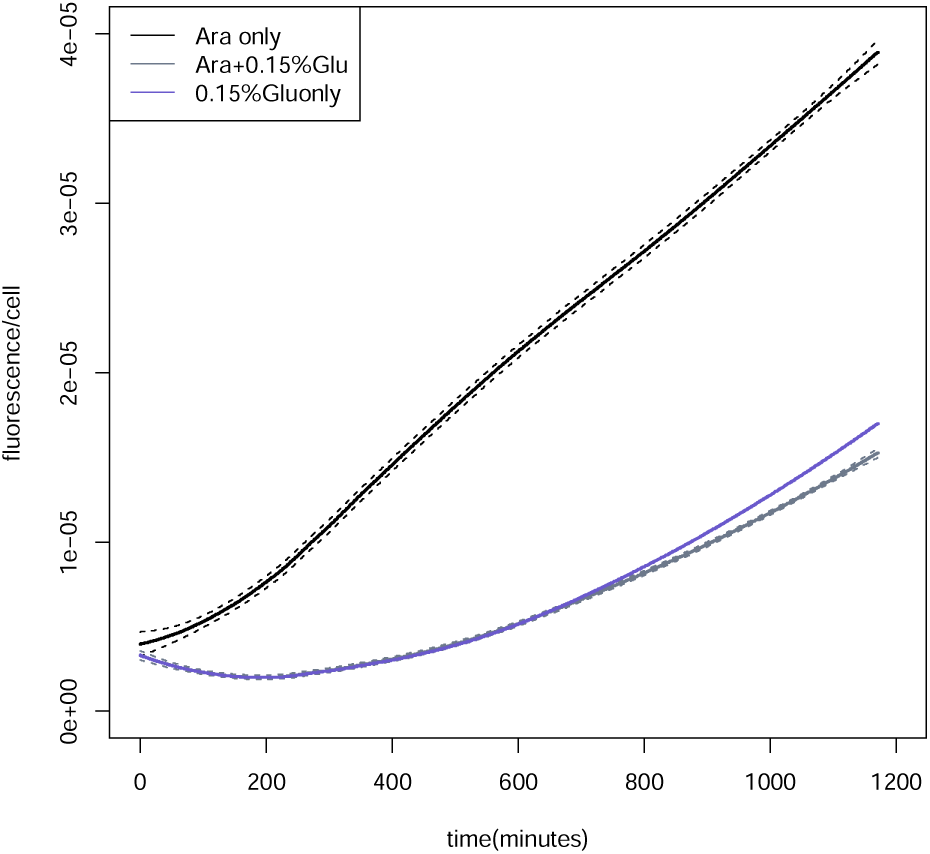
- Removal of glucose affects ROS production in *E. coli*. Fluorescence per cell was measured over 1200 minutes for TOP10 cells in the absence of the M.SssI plasmid. 0.15% glucose is sufficient to repress ROS levels, demonstrating that it is removal of glucose rather than addition of arabinose that causes ROS accumulation.

**Supplemental Figure 6.**
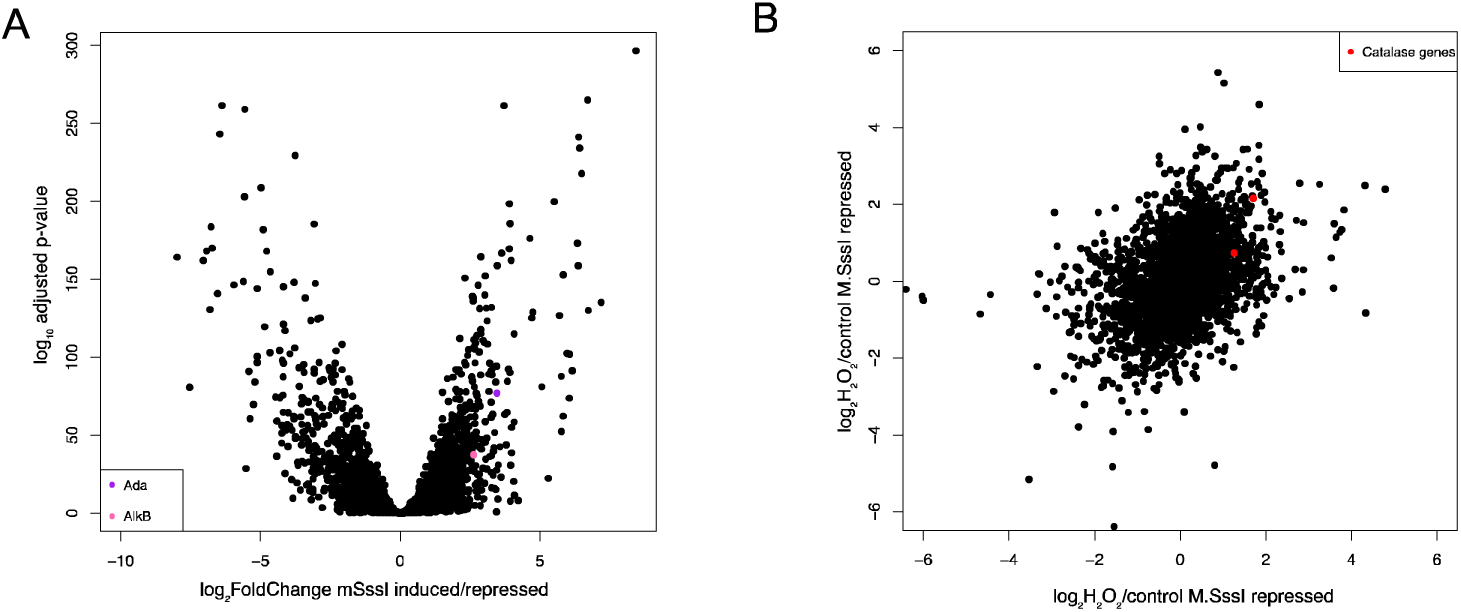
- Transcriptomics analysis. A shows a volcano plot of genome wide transcript level changes measured by RNAseq comparing M.SssI expressing cells (arabinose) to M.SssI repressed cells (glucose). The x axis shows log_2_ fold change and the y axis shows the Benjamani Hochberg adjusted p-value calculated through DESeq2. *alkB* and *ada* are highlighted. B shows the log2 fold change in transcript levels upon H2O2 treatment of cells in which M.SssI is repressed (x axis) compared to cells where M.SssI is expressed (y axis). Catalase genes *katG* and *katE* are highlighted.

**Supplemental Figure 7.**
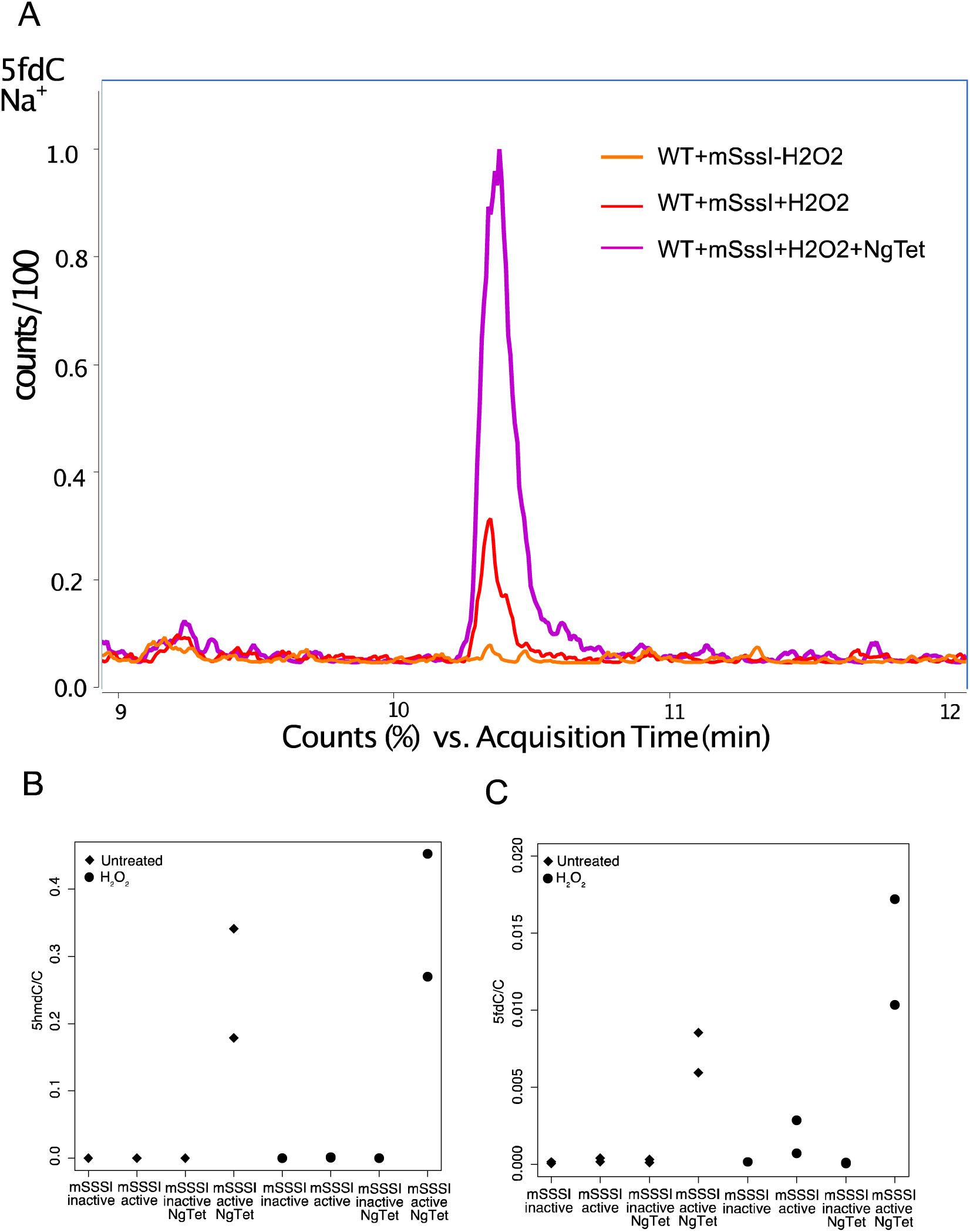
- TET expression induces 5hmC and 5fmC in E. coli Top: example trace showing 5fC levels in cells expressing mSssI either with or without H2O2 and in the presence of Tet from *N. gruberii*. Bottom: quantification of 5hmC (left) and 5fC (rights) demonstrating induction of 5hmC and 5fC by Ng Tet expression, and induction of 5hmC and 5fmC at low levels by H_2_O_2_ alone.

## Supplemental Material

SupTables1.xls Spreadsheets containing all the raw data underlying figures 1,2,3,5 and 6.

SupTables2.xls Spreadsheets containing raw and processed data for figure 4 (ROS measurements)

SupTable3.xls Mass spectrometry data

SupFile4.Rdata Transcriptomics data raw counts

